# Magnetic particle imaging reveals that iron-labeled extracellular vesicles accumulate in brains of mice with metastases

**DOI:** 10.1101/2024.03.12.584146

**Authors:** Victoria A Toomajian, Anthony Tundo, Evran E Ural, Emily M Greeson, Christopher H Contag, Ashley V Makela

**Author notes:** Corresponding author: Ashley V Makela.

## Abstract

The incidence of breast cancer remains high worldwide and is associated with a significant risk of metastasis to the brain that can be fatal; this is due, in part, to the inability of therapeutics to cross the blood brain barrier (BBB). Extracellular vesicles (EVs) have been found to cross the BBB and further, have been used to deliver drugs to tumors. EVs from different cell types appear to have different patterns of accumulation and retention as well as efficiency of bioactive cargo delivery to recipient cells in the body. Engineering EVs as delivery tools to treat brain metastases, therefore, will require an understanding of the timing of EV accumulation, and their localization relative to metastatic sites. Magnetic particle imaging (MPI) is a sensitive and quantitative imaging method that directly detects superparamagnetic iron. Here, we demonstrate MPI as a novel tool to characterize EV biodistribution in metastatic disease after labeling EVs with superparamagnetic iron oxide (SPIO) nanoparticles. Iron-labeled EVs (FeEVs) were collected from iron-labeled parental primary 4T1 tumor cells and brain-seeking 4T1BR5 cells, followed by injection into mice with orthotopic tumors or brain metastases. MPI quantification revealed that FeEVs were retained for longer in orthotopic mammary carcinomas compared to SPIOs. MPI signal due to iron could only be detected in brains of mice bearing brain metastases after injection of FeEVs, but not SPIOs, or FeEVs when mice did not have brain metastases. These findings indicate the potential use of EVs as a therapeutic delivery tool in primary and metastatic tumors.

**TOC Graphic:** 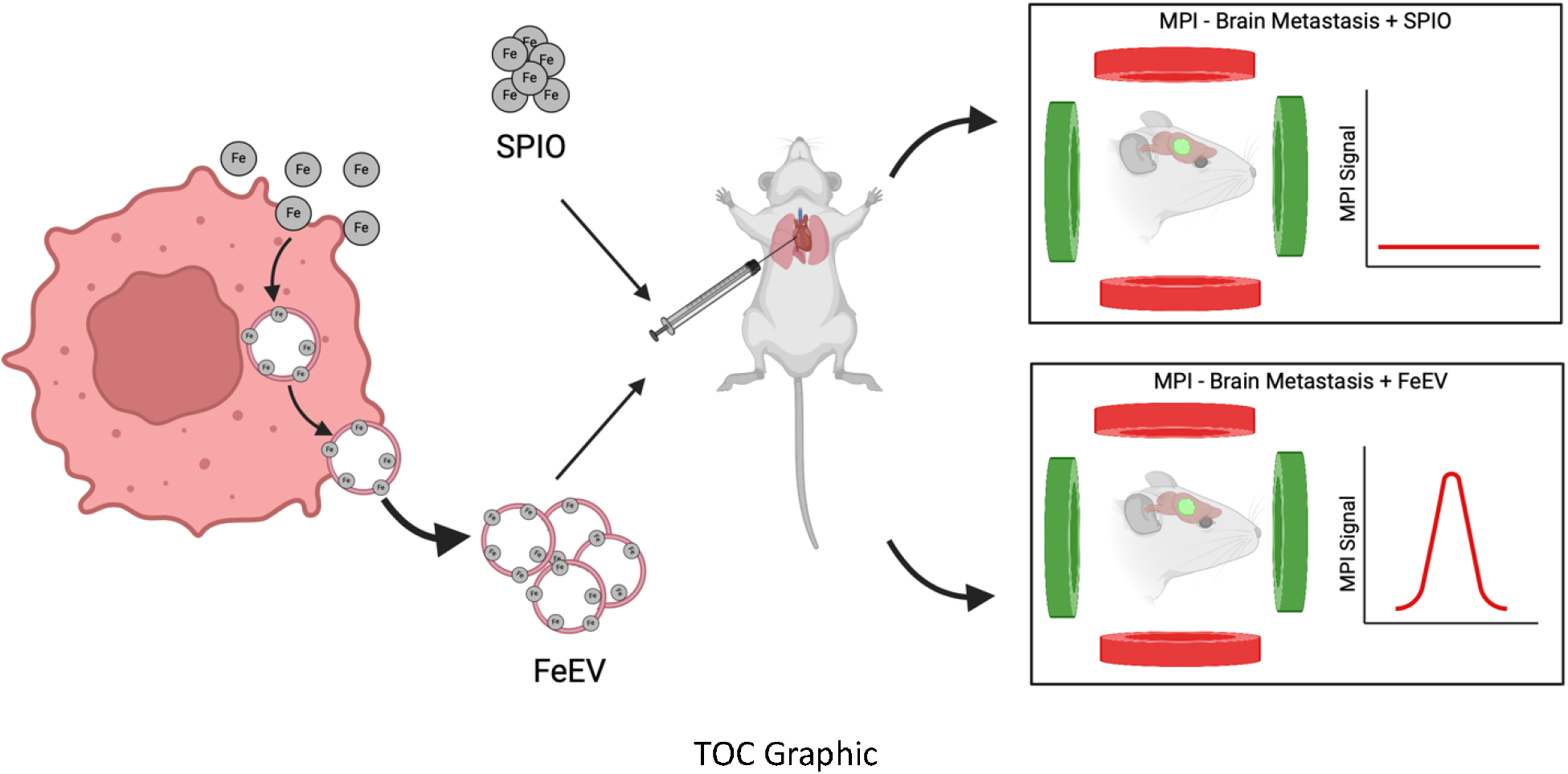

## Introduction

Among all new cases of cancer worldwide in 2022, the incidence of breast cancer was the highest in females at 23.8%^1^. Although there have been advances in treating the primary tumor, metastases continue to be the leading cause of death among these patients^2^. In particular, up to 30% of breast cancer patients have brain metastases^2,3^ and the prognosis for these patients is poor with a survival rate of only 20% at 1 year^3–6^.

Treatment of brain metastasis includes radiotherapy and surgery, and use of these options is dependent on the amount of metastatic burden in the brain. However, these have limited survival benefit and can result in a reduced quality of life^7,8^. Systemic treatments such as cytotoxic chemotherapy may be used, but these therapies have difficulty reaching the tumor due to the presence of the blood-brain-barrier (BBB) which is formed by endothelial cells and tight junctions^9,10^. When the BBB remains intact, the use of a systemic treatment will not reach the target and not elicit any therapeutic outcome. However, the BBB can be disrupted in a heterogenous manner with the formation of a tumor, resulting in what is known as the blood-tumor-barrier (BTB)^9,11^. This heterogenous permeability causes changes in accumulation of chemotherapies; this has been observed in experimental models of brain metastases where there has been evidence of impairment of the BTB, although some barrier functions remain intact, limiting the accumulation of drugs to such low amounts which do not elicit any therapeutic effect^12,13^. A strategy for improving delivery of therapeutics across these barriers is needed to effectively combat brain metastases.

One possibility is to use naturally derived particles with lipid membranes to deliver therapeutics across the BBB or improve delivery across the BTB. Extracellular vesicles (EVs) are small particles naturally released from cells that carry and deliver bioactive molecules as a method of cell-to-cell communication, and EVs from various types of cells have been shown to target tumors and cross the BBB^14,15^. EVs can be classified by mechanism of biogenesis, size, function, or composition; examples of EVs classified by biogenesis include exosomes, microvesicles, and apoptotic bodies^16^. Exosomes are formed by internal budding within a multivesicular body, and range in size from 40 to 150 nm in diameter. Microvesicles are formed by an external budding of the cell membrane and are typically 100 nm to 1 µm in diameter. Apoptotic bodies, the largest class of vesicles, are released from dead and dying cells and can be up to several microns in diameter. Because of the overlapping size range of microvesicles and exosomes, EVs are often characterized as enriched populations of small EVs and large EVs rather than pure populations^16^. Because EVs can circulate in the bloodstream for a prolonged period of time, can cross the BBB^14,15,17–22^, and are manipulable as carriers of drugs, nucleic acids, or nanoparticles^19–31^, there is vast potential to engineer EVs as imaging or delivery tools to target brain metastases. Previous studies have labeled EVs with superparamagnetic iron oxide (SPIO) nanoparticles for *in vivo* tracking using magnetic resonance imaging (MRI)^26–28,32,33^ and to a lesser extent, magnetic particle imaging (MPI)^34^. Additionally, EVs derived from tumor cells have been shown to improve delivery to parental tumor cells *in vivo*^35–37^. A recent study has used photoacoustic imaging to detect Prussian blue nanoparticles within an orthotopic brain tumor model in mice^38^. Coating the particles with glioma U-87 derived EVs facilitated the delivery of the Prussian blue nanoparticles into the tumor region within the brain.

An imaging technique which could facilitate the improvement of EV delivery across the BBB or BTB would offer significant benefits for treating patients suffering from brain metastases. Identifying localization and accumulation in the region of interest would provide confirmation of successful delivery and assess retention over time. MPI is a sensitive and quantitative imaging method that detects the magnetic properties of SPIOs directly^39–41^ and offers the opportunity to assess the biodistribution of EVs associated with SPIOs. MPI benefits from essentially no background magnetic signals in tissue, with signals originating only from SPIOs. In addition, there is no loss of signal due to mammalian tissues and thus the depth of particles within the body does not adversely affect quantitative imaging. MPI has been used for cancer detection^39,42–45^ and has been used to track engineered EVs to primary tumors^34^ in animal models. In addition to this, other applications of MPI have included vascular imaging^46–50^, inflammation^51,52^, therapeutic studies^53,54^ and cell tracking^54–60^. Further, groups are working towards improved nanoparticles tailored for MPI^61–65^, improvements in MPI hardware and acquisition^66,67^ and image analysis^68,69^. Other imaging methods such as *in vivo* bioluminescence imaging (BLI) can be used as a complementary imaging modality in rodent models to associate or correlate MPI signals with other biological processes such as sites of metastatic lesions^70^.

In this study, we labeled murine breast cancer cells and a brain seeking derivative cell line (4T1 and 4T1BR5, respectively) with SPIOs, which led to production of iron-labeled EVs (FeEVs) in the culture medium. The FeEVs were tracked using MPI to monitor their retention in a primary breast tumor or accumulation in the heads of mice which had brain metastasis.

## Results and Discussion

### Labeling Cells with Iron Leads to Production of Iron-Labeled EVs

Co-incubation of 4T1-fLuc2 (4T1L2) or 4T1BR5-fLuc2/GFP (4T1BR5-L2G) cells in culture with SPIO nanoparticles resulted in cell secretion of iron-labeled EVs (FeEVs) into the culture medium. This method of protamine sulfate/heparin assemblies to aid in Synomag-D internalization into 4T1BR5 cells has been performed previously, with no changes in viability of cells up to 3 days post labeling^42^. Upon isolation from conditioned medium via differential centrifugation, the SPIOs were found to be associated with the cell-secreted EVs, as confirmed using transmission electron microscopy (TEM) visualizing the EV membranes (stained with uranyl acetate for contrast) and our dextran-coated iron nanoparticles. Free iron particles (red arrowheads) and iron particles associated with EVs (red arrows) were observed in the samples (**Figure 1a,c**; 4T1L2 and 4T1BR5-L2G, respectively). Images showed iron to be associated with the EV membrane; it did not appear that the EV membranes encapsulated the SPIOs. After washing, a large amount of what appeared to be free iron particles were still present in the preparations. The FeEVs and EVs from cells which were not cultured with iron (**Figure 1b,d;** 4T1L2 and 4T1BR5-L2G, respectively) appeared to have a similar size and morphology, both appearing round (yellow arrowheads) and often having the classical cup shape (yellow arrow).

**Figure 1.**
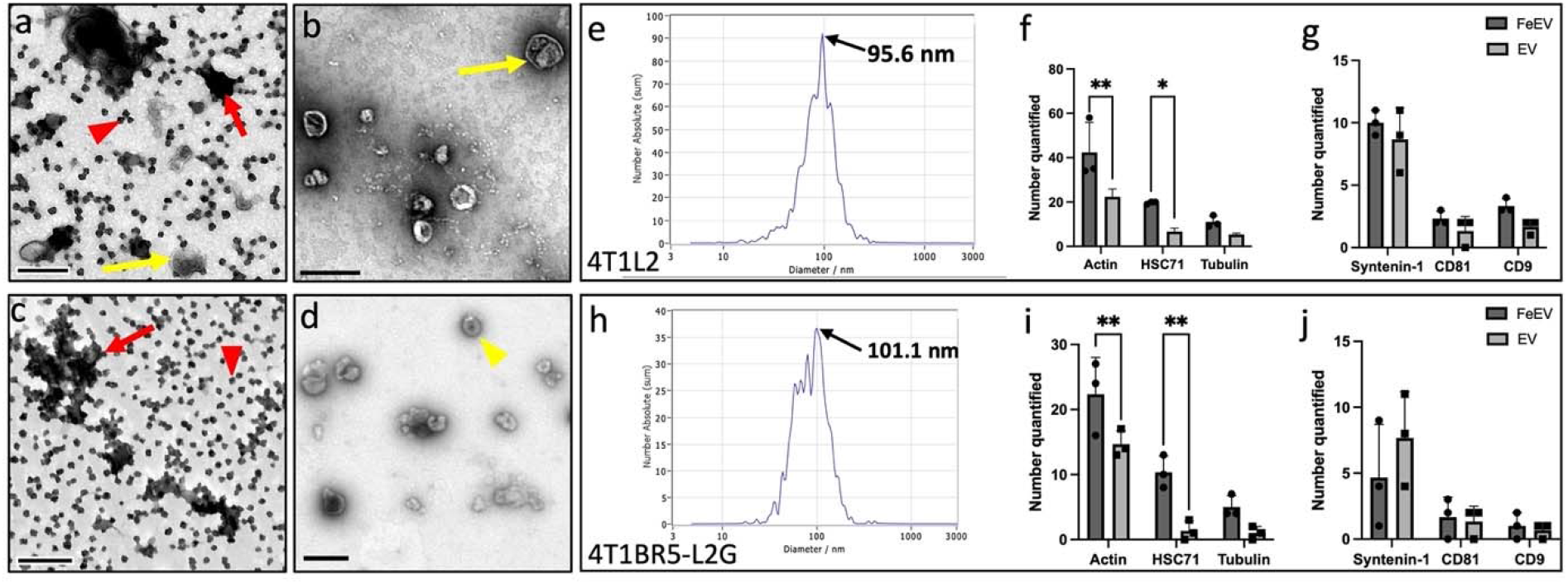
Iron-labeled extracellular vesicle characterization. Iron-labeled EVs (FeEVs) (**a**,**c**, 4T1L2 and 4T1BR5-L2G, respectively) and unlabeled EVs (**b**,**d**, 4T1L2 and 4T1BR5-L2G, respectively) were visualized via transmission electron microscopy (TEM). Both free iron (red arrowheads) and EVs associated with iron (red arrows) can be seen in the FeEV samples. FeEVs and EVs both display the classical cup shape (yellow arrows) and round shape (yellow arrowhead). Nanoparticle tracking analysis determined peak FeEV size in 4T1L2 (**e**) and 4T1BR5-L2G (**h**) cells. Abundance of cytosolic and membrane proteins typically associated with EVs were quantified in 4T1L2 (**f**,**g**) and 4T1BR5-L2G (**i**,**j**) EVs and FeEVs as detected by mass spectrometry. Scale bars = 250 nm. \**p*<.05, \*\**p*<.01

Enriched FeEV samples were characterized by examining the particle size and composition with regard to iron content and proteins. Iron content per EV was quantified by imaging different masses of Synomag-D by MPI (**Figure S1a**). The linear relationship between iron mass and MPI signal (**Figure S1b**, R^2^=.996), allows for a quantitative measure of iron (**Table S1**) in the FeEV pellets imaged by MPI (**Figure S1c**). The average peak size of the 4T1L2 cell-derived FeEVs was 98.8 +/-4.98 nm (**Figure 1e**, representative image), and 98.43 +/-13.79 nm in 4T1BR5-L2G cell-derived FeEVs (**Figure 1h**, representative image). These sizes are within the average size of small EVs, typically defined as 30-150 nm^71^, but larger than the 70 nm native SPIO particles. The FeEVs contained similar iron loading, with 0.57 fg iron per EV (Fe/EV) and 0.51 fg Fe/EV in 4T1L2 and 4T1BR5-L2G FeEVs, respectively. Based on a lower MPI detection limit of ∼313 ng of Synomag-D (in 1 µl), using our standard imaging parameters (**Figure S1a**), we could expect a sensitivity of 5.52-7.42×10^8^ FeEVs, using the average Fe/EV for 4T1BGL, 4T1L2 and 4T1BR5 derived FeEVs. However, the volume in which the FeEVs accumulate may vary, resulting in a sensitivity higher or lower than our prediction.

Mass spectrometry analysis of FeEVs and EVs derived from 4T1L2 (**Figure 1f, g**) and 4T1BR5-L2G (**Figure 1i, j**) cells showed expression of ^71^EV-associated cytoskeletal proteins actin and tubulin α-1A, known to be promiscuously incorporated in EVs^71^. Further, heat shock cognate 71 (HSC71), tetraspanins CD81 and CD9, and Syntenin-1 were also identified in the samples. FeEVs contained significantly more actin and HSC71 in both 4T1L2 (*p*=.001 and *p*=.02, respectively) and 4T1BR5-L2G (*p*=.007 and *p*=.002, respectively) when compared to EVs without iron. Actin and HSC71 have been identified as phagosomal proteins^72,73^, so the increased expression observed in FeEVs may be due to phagocytic activity in cells upon coincubation with iron, and promiscuous incorporation into the secreted FeEVs.

There was no difference in tubulin α-1A expression between FeEVs and EVs for either 4T1L2 or 4T1BR5-L2G cells (*p*=.26 and *p*=.113, respectively). No differences were found in expression of syntenin-1, CD81, and CD9 between FeEVs and EVs in 4T1L2 (*p*=.223, *p*=.354 and *p*=.134, respectively) and 4T1BR5-L2G cells (*p*=.147, *p*=.866 and *p*=.866, respectively). Overall, cellular incorporation of iron into EVs did not affect expression of typical EV markers involved in EV biogenesis and release, which has been observed previously^74^.

Mass spectrometry analysis was confirmed via Western blot analysis of FeEVs, EVs and cell lysate samples derived from 4T1BR5-L2G cells, which all showed expression of Alix and Flotillin-1^71^ (**Figure S2**). Super resolution microscopy confirmed the presence of tetraspanins CD81 (magenta), CD9 (yellow) and CD63 (cyan) in samples of FeEVs derived from 4T1BR5-L2G cells (**Figure S3**). Aside from being typical EV markers, tetraspanins are also implicated in a number of processes^75,76^. Although recently the tetraspanins CD63 and CD9 have been shown to not be required for EV uptake and content delivery^77^, we did observe the tetraspanin CD81 and flotillin-1, both of which have been implicated in EV uptake^75,76,78–81^.

4T1 extracellular vesicles have been studied and characterized previously^82–84^, and there is evidence of preferential homing of EVs to a matched parent cancer cell line *in vitro*^36,85,86^. The similarities in composition of the EV (*i.e*., lipid composition and surface receptors) and the matched parent cell line increase the propensity of fusion and internalization^84^. Further, EVs do not exhibit any cytotoxicity when incubated with 4T1 cells in culture^87^, highlighting their biocompatibility. Improved internalization of SPIO into 4T1BR5-L2G cells was seen when the SPIO was associated with EVs (FeEVs) versus the SPIO alone (**Figure 2**). All cells, identified by DAPI (blue) and GFP (green), had PKH26 stained FeEVs (membrane stain, yellow and SPIO far-red fluorescent, magenta), to varying extents (**Figure 2a**). In some instances, the PKH26 and SPIO fluorescence (yellow and magenta, respectively), were located in the same spatial location (thick arrow, **Figure 2b**), suggesting the association of PKH26-stained EVs and the SPIO. PKH26 without SPIO signal was found in other locations (arrowhead, **Figure 2b**), suggesting the presence of unlabeled EVs. When cells were incubated with SPIO only (**Figure 2c**), there was minimal amounts of SPIO found within the cells. Both extracellular SPIO was found (**Figure 2d**, dashed line) and SPIO without PKH26 signal within cells (**Figure 2d**, thin arrow).

**Figure 2.**
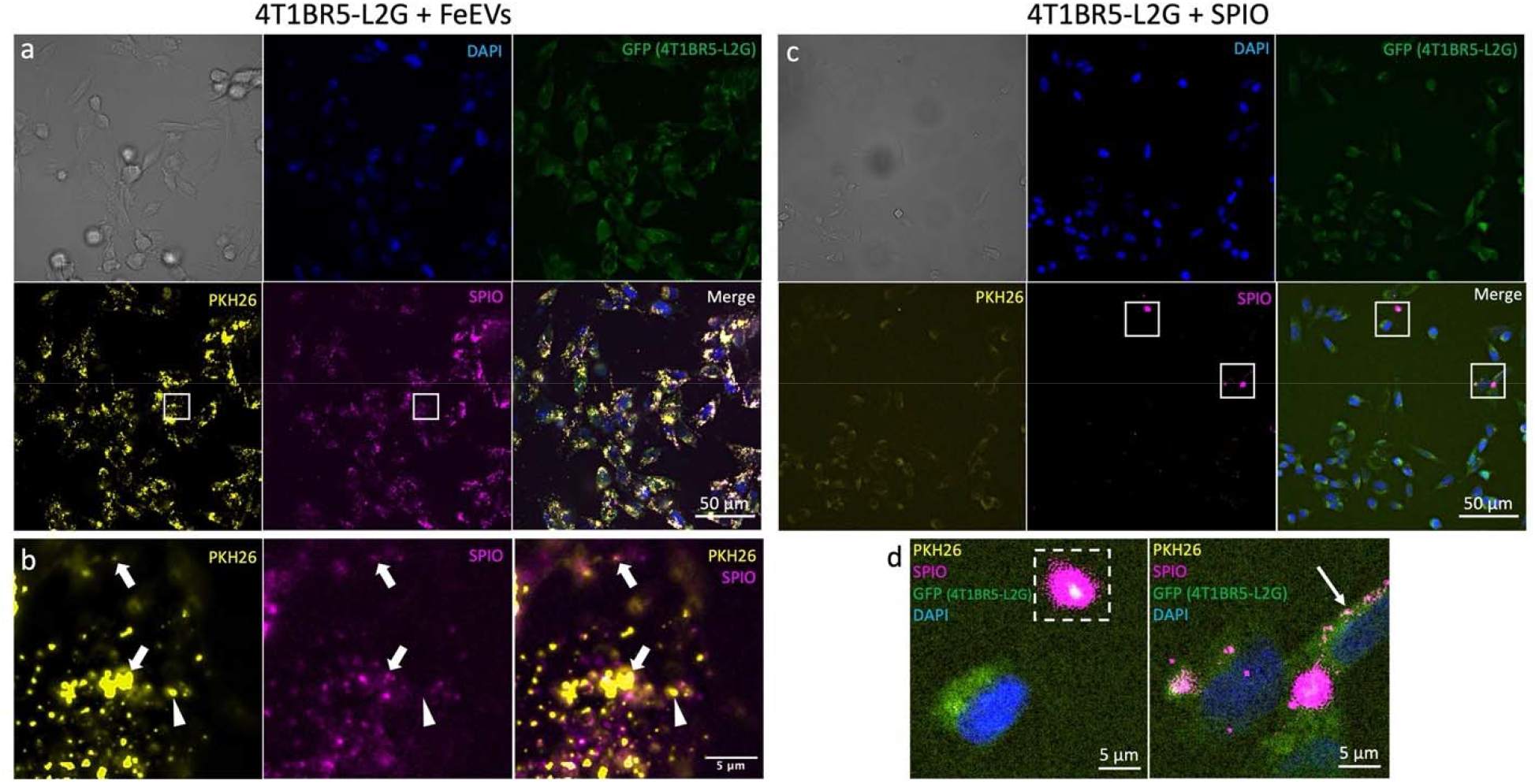
SPIO associated with EVs (FeEVs) accumulate more than SPIO alone in 4T1BR5-L2G cells in culture. After 24 hours incubation, there was more SPIO in 4T1BR5-L2G cells when introduced as FeEVs (**a**) versus SPIO alone (**c**). The 4T1BR5-L2G cells are identified by blue (DAPI = nuclei) and green (GFP). The SPIO was tagged with a far-red fluorophore (magenta) and FeEVs or SPIO were dyed with PKH26 (yellow) to identify membranes. In the samples which were incubated with FeEVs (**b**, zoomed in from area identified in **a**), there are both regions with PKH26 and far-red SPIO fluorescence in the same spatial location (thick arrows, **b**), as well as PKH26 without far-red SPIO signal (arrowheads, **b**). In the samples which were incubated with SPIO alone, there was extracellular far-red SPIO signal noted (dashed outline, **d**) and far-red SPIO appearing within the cells (thin arrow, d).

### Using MPI to understand the biodistribution of FeEVs and SPIOs

FeEVs and SPIOs were injected intravenously (i.v.) into healthy and 4T1 orthotopic tumor-bearing mice to understand the kinetics of FeEVs and SPIO biodistribution over time *in vivo*. When 4T1-derived FeEVs or equal amount of iron were administered i.v. into healthy (FeEVs: 1.45×10^10^, SPIO: 8 µg) or tumor-bearing mice (FeEVs: 1.7×10^10^), MPI detected iron in the liver (**Figure S4**) but not in primary tumors. The high accumulation in the liver has resulted in challenges identifying and quantifying iron in regions of close proximity, by MPI^88^. Here, this would result in difficulties visualizing the smaller amounts of iron which may have accumulated in the tumor.

### Primary mammary fat pad tumors retain FeEVs more than free iron

FeEVs and SPIOs were injected intratumorally (i.t.) into 4T1 orthotopic tumor-bearing mice to compare iron retention over time *in vivo*. The i.t. injection of FeEVs (1.7×10^10^ FeEVs) or equal amount of iron delivered via SPIOs (9.6 µg) resulted in different retention dynamics (**Figure 3**). Localization of the iron was detected via MPI immediately post injection of FeEVs and SPIOs (**Figure 3a,b**). In tumors which received FeEVs injections, a higher percentage of iron was quantified from the injected dose (93.4%) immediately (0-h) after injection as compared to tumors injected with SPIO (63.6%, *p*=.009, **Figure 3c**). There was a significant decrease in the amount of iron detected 24-hours (h) after the injection of FeEVs into tumors (69.2% of iron quantified at 0-h, *p*=.026), but the amount of iron detected at 72-h post FeEV injection did not decrease further (53% of iron quantified at 0-h, *p*=.06). However, there was a large decrease in the amount of iron quantified 24-h after SPIO injection into the tumors (6.8% of iron detected at 0-h, *p*=.002), with no further difference in the amount of iron detected at 72-h post injection (*p*=.373, compared to 24-h), with 14.9% of iron quantified at 0-h present. Immediately post injection, significantly more iron detected in tumors injected with FeEVs suggests that there is rapid clearance of free SPIO from the tumor, whereas there was an initial retention of iron associated with EVs. At 24-h post-injection there is a decrease in iron quantified in both FeEV and SPIO-treated groups. However, the tumors injected with FeEVs or SPIO had no significant difference in iron quantified at 24-h or 72-h, suggesting the vast majority of SPIO clearance from the tumor occurs within the first 24-h of injection, while tumor retention is promoted in iron associated with EV membranes.

**Figure 3.**
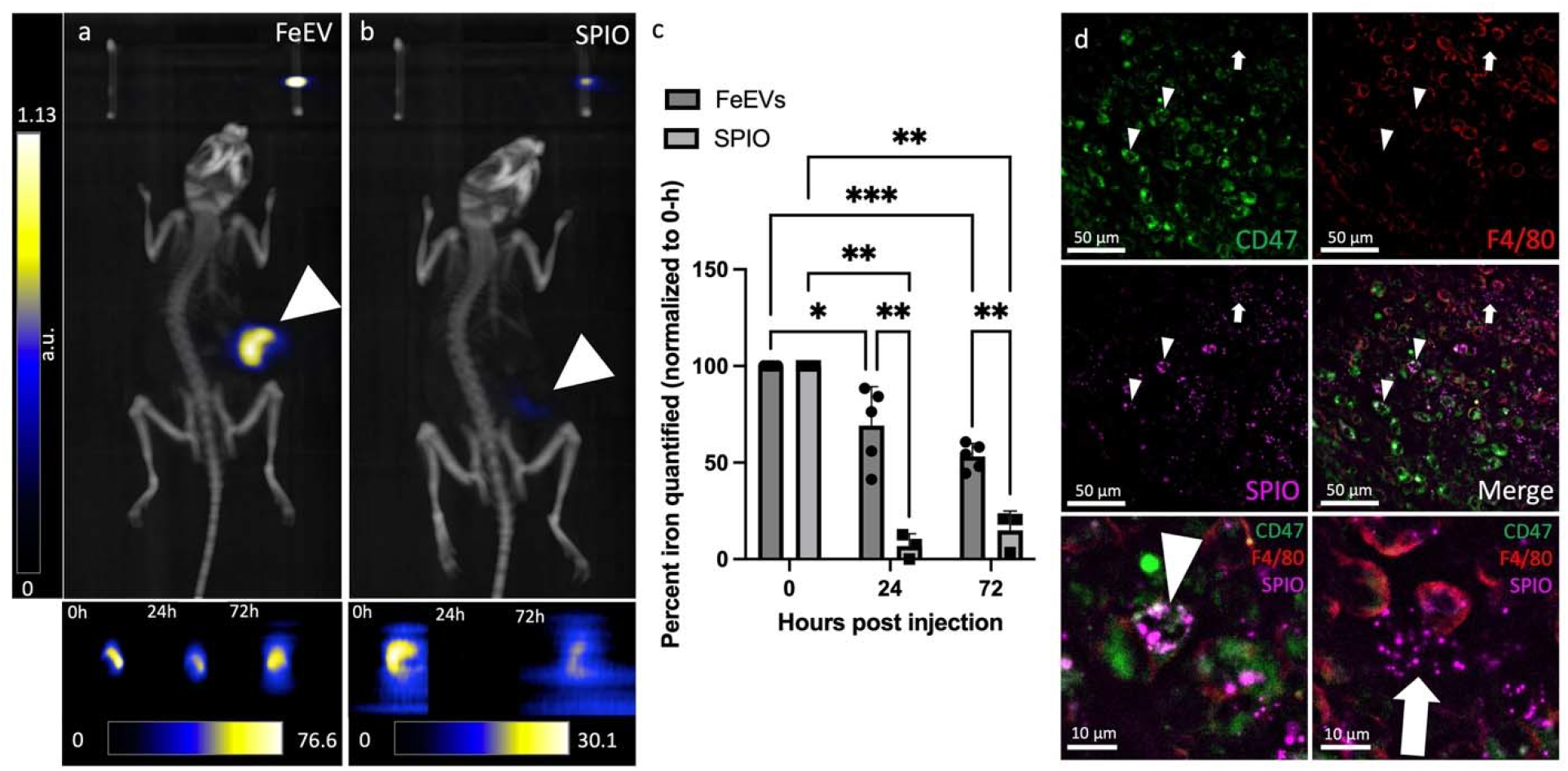
Increased retention of FeEVs in the location of the primary tumor compared to SPIO. Magnetic particle imaging and microCT images (**a**,**b**, representative images, 72-h post injection) were overlayed to localize iron signal after intratumoral injection of FeEVs (**a**) or SPIO (**b**) with cropped 2D MPI images of the tumor at 0, 24 and 72h located below. Location of tumors noted by white arrowhead. Tumor iron quantification (**c**) at 0, 24 and 72-h post injection for FeEV- and SPIO-injected tumors (normalized to iron quantified at 0-h). Fluorescence microscopy (**d**) identifies SPIO (far-red, magenta), CD47 tumor cells (PE, green) and F4/80+ cells (AF647, red). SPIO and CD47+ cells in the same spatial location (arrowheads) and SPIO and F4/80+ cells within the same region (arrow). \**p*<.05, \*\**p*<.01, \*\*\**p*<.0001

Histology of tumors excised at 72-h post injection revealed the localization of FeEVs. No iron was detected in the mammary fat pad (MFP) tumors that had been injected with SPIOs, corresponding to the very low amount of iron detected by MPI. However, within the tumors injected with FeEVs, iron (magenta) was found inside CD47+ tumor cells (green; **Figure 3d**, white arrow head) but not F4/80+ cells (red; **Figure 3d**, white arrow), indicating that cargos associated with FeEVs could be delivered to tumor cells. F4/80 serves as a common macrophage surface marker, although it can also be used to define dendritic cells^89^. Further, 4T1 tumors have been shown to contain a high number of tumor-associated macrophages (TAMs)^90,91^, validating our findings. While CD47+ tumor cells showed SPIO uptake with a cell-like shape, F4/80+ cells did not appear to contain SPIO; rather, it was located near these cells. It is unlikely the tumor cells internalized SPIO alone, as SPIOs administered *in vivo* are typically only found in macrophages^88,90,92^. This demonstrates that FeEVs can communicate or deliver cargo to tumor cells, which suggests the likely reason the iron associated with EVs was detected by MPI in greater quantity and for longer than SPIOs.

EVs derived from 4T1 cells have been used previously *in vivo*, to identify localization to targeted metastatic sites (lungs, liver and spine)^83^, and their targeted delivery of cargo to parental 4T1 tumors^82^. These studies, along with our immunocytochemistry, suggest that cellular origin has an influence on the localization of the EVs after administration *in vivo*.

### Association of iron with EVs allows iron to accumulate in the heads of mice with brain metastasis

Experimental brain metastases were initiated by administering 4T1BR5-L2G cells into the left ventricle of the heart (intracardiac, i.c.). To confirm successful delivery, mice were imaged using BLI to confirm tumor cell localization in the brain (**Figure 4a**) and continued to observe growth of brain metastases. Following i.c. administration of FeEVs (6.97×10^10^, average of 33.64 +/-5.44 µg Fe), the majority of iron (as determined by MPI signal) was found in the liver (**Figure 4b**) both in mice burdened with brain metastasis (93% of injected) or healthy controls (102% of injected). Thus, the liver still filters much of the FeEVs and SPIOs, despite injection into the left ventricle.

**Figure 4.**
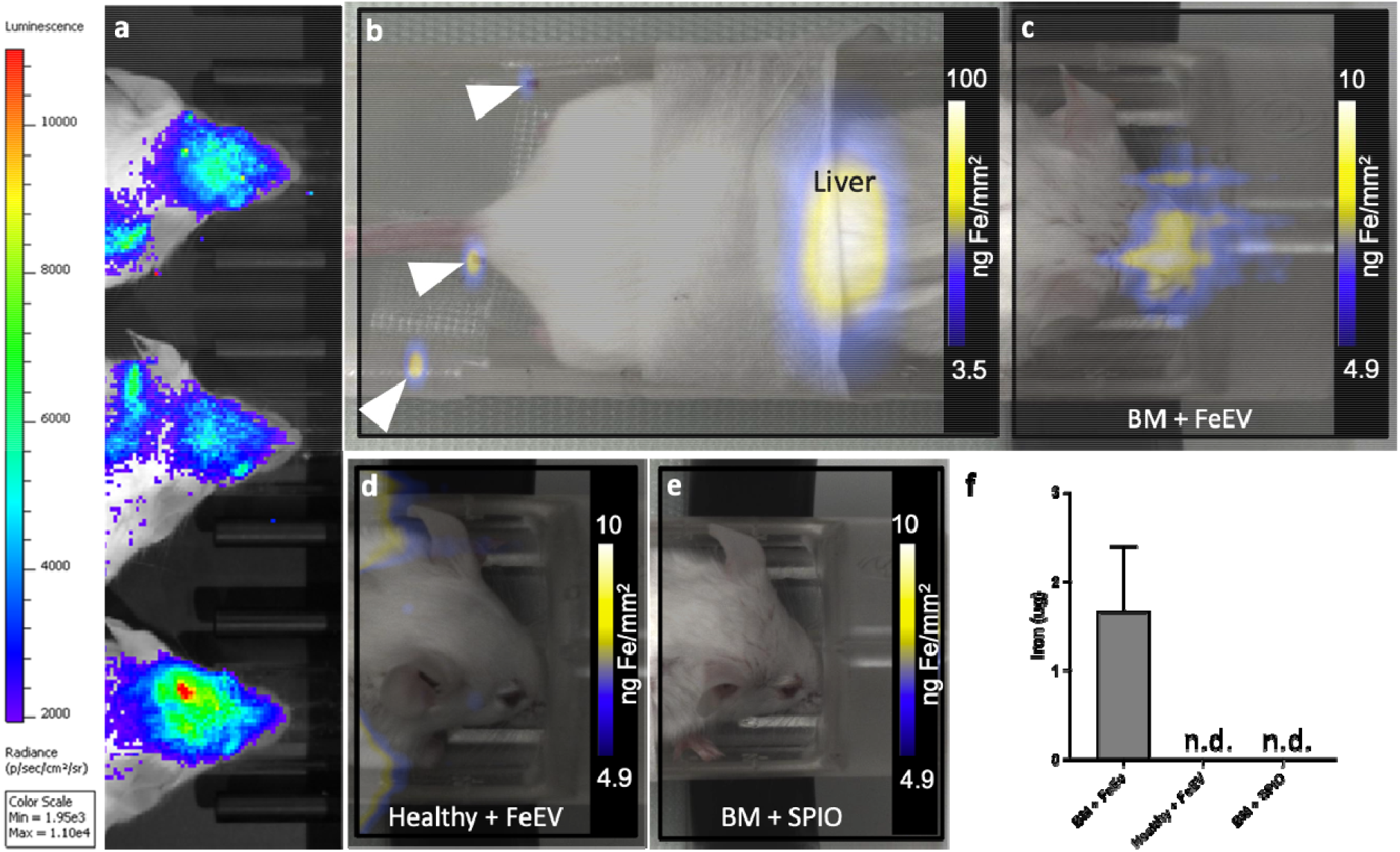
Magnetic particle imaging detects and quantifies iron in heads of mice. Bioluminescence imaging (**a)** was used to confirm brain metastasis in the heads of mice which received intracardiac administration of 4T1BR5-L2G cells. MPI and brightfield images were overlayed to localize MPI signal. MPI signal was located in the liver (representative image, **b)** in all groups. White arrowheads note location of iron fiducials. MPI signal was only located in the head of mice with brain metastasis which were injected with FeEVs (**c**, BM + FeEV). There was no signal located in mice which did not have brain metastasis and received FeEVs (**d**, Healthy + FeEVs) or mice which had brain metastasis and received SPIO (**e**, BM + SPIO). MPI was used to quantify iron in the heads of mice from each group (**f)**. n.d. = **n**ot detected.

MPI signal due to the presence of iron was detected in the heads of 10/11 mice with brain metastases post-FeEV injection, with iron amounts averaging 1.67 +/-0.72 µg Fe (**Figure 4c**), amounting to 4.96% of the injected dose. Furthermore, no MPI signal was detected in the heads of healthy mice i.c. injected with FeEVs, or in the heads of mice with brain metastases i.c. injected with SPIOs (**Figure 4d**,**e**). This demonstrated improved FeEV targeting to brain metastasis and delivery of cargo, implying that FeEVs have potential for future diagnostic imaging and therapeutic targeting.

EVs have been used to deliver payloads to the brain in mice through intranasal administration^21,22,93–95^ and systemic injection^20,26,27^. In addition to their ability to cross the BBB, EV use is favorable because of low immunogenicity, biodegradability, and their non-toxic nature. Further, they can carry cargo, such as iron particles in this study and others^26,28,30,33,34,96,97^, as well as therapeutics^19,20,22–25,27,94,95^. The mechanisms in which EVs cross the BBB is not fully understood; however, some possible routes which have been explored are micropinocytosis, clathrin-dependent endocytosis and caveolae-dependent endocytosis^17,18,98,99^.

### Histology reveals iron accumulation in brains

Brain sections from FeEV and SPIO-injected mice with brain metastases and FeEV-injected healthy mice were stained with Perls Prussian Blue (PPB; blue = iron, pseudocolored magenta in overlay) to visualize iron and DAPI to visualize nuclei. Within brain sections from mice with metastasis injected with FeEVs, we found iron associated with brain metastasis (as determined by changes in tissue architecture with increased and disorganized nuclei by DAPI)^100^, as well as within regions without an apparent brain metastasis (**Figure 5a**). There were also brain metastasis which did not have any iron associated with them (**Figure 5a**). In sections derived from brains of healthy mice injected with FeEVs, there was iron found in lesser amounts (**Figure 5b**). Further, the group with brain metastasis that received SPIO also had iron found in the brain sections, but no iron was found associated with brain metastasis (**Figure 5c**), and the iron identified was in lesser amounts. There was more apparent iron seen within the brain sections of mice with brain metastasis injected with FeEVs; the localization of the iron also appeared more intentional, either within the metastasis or surrounding it. Comparatively, iron in mice with brain metastasis injected with SPIOs and healthy mice injected with FeEVs were in lesser amounts, correlating with the undetectable iron levels in MPI.

**Figure 5.**
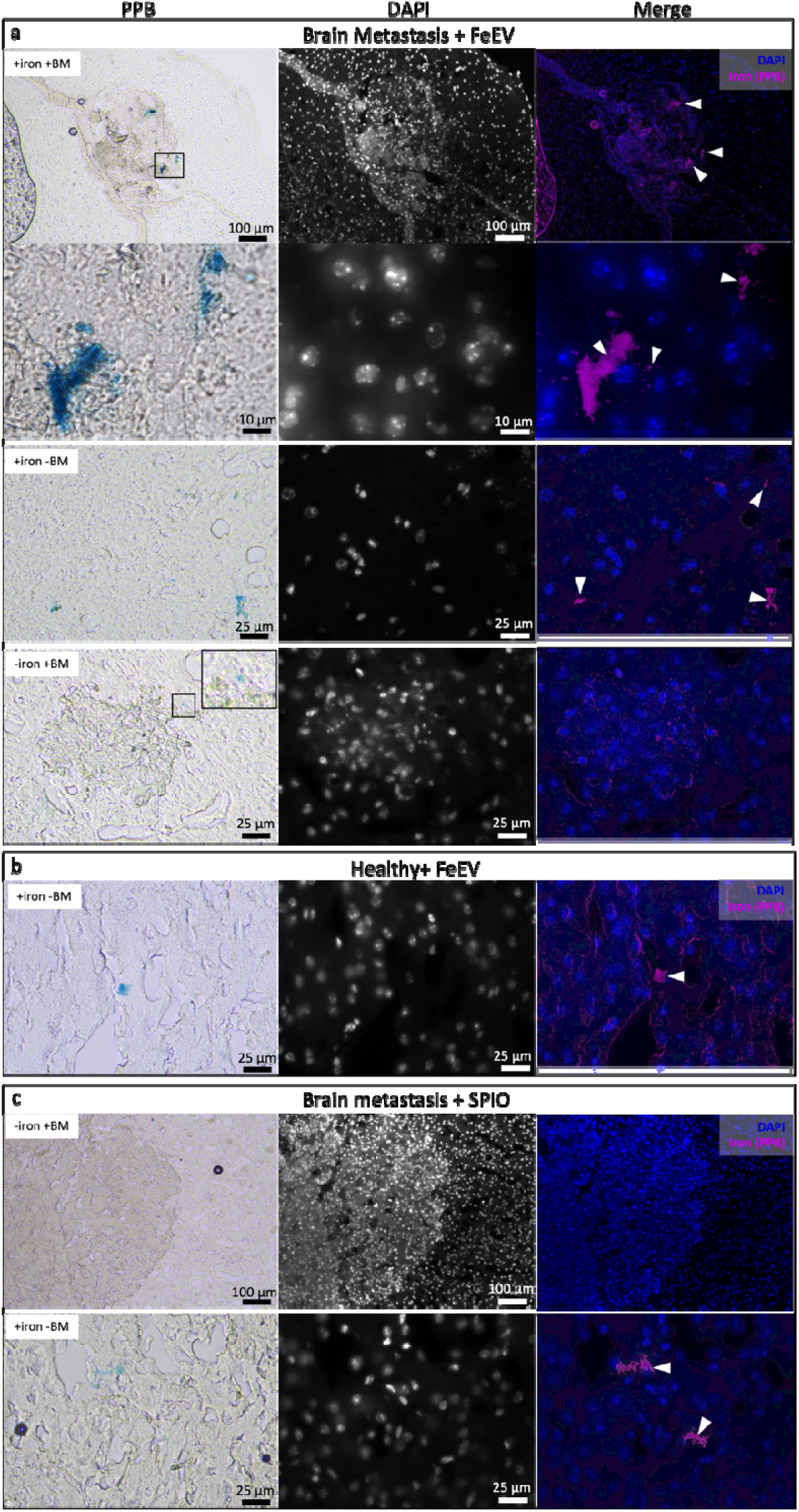
Histology locates iron present in brain sections. Sections were stained with Perls Prussian Blue for iron (left column) and DAPI to visualize nuclei (middle column). Iron was pseudocolored magenta to overlay with DAPI (blue, right column). Sections from the group which had brain metastasis and received FeEVs (**a)** had regions where there was iron within metastasis, iron outside of metastasis and metastasis not associated with iron. Mice which did not have metastasis and received iron (**b)** did have some iron visualized. Mice which had metastasis and received iron (SPIO) had metastasis visualized without iron and iron within regions of no metastases.

Experimental brain metastasis models, including the 4T1BR5 model used in this study, have shown variable effects on the BTB. Although the BTB is impaired in the majority of these metastases, fewer than 10% exhibit leakiness which allows for enough drug to enter to elicit therapeutic effects^12,13^. BTB impairment could explain the presence of iron identified by PPB, with passive delivery of the iron whereas an increase in iron accumulation when associated with EVs (FeEVs) can be considered an improvement in delivery, independent of BBB/BTB status. Therefore, there is still a need to develop ways to improve the delivery both when the BBB is intact, or even when there is BTB heterogeneity in the tumors.

## Conclusions

Iron-labeled EVs have been imaged previously using MRI^27–29,32,33^ and MPI^34^. Methods used to label EVs have been through direct labeling, or indirect labeling where the parental cell is co-incubated with the iron and is allowed to take up the nanoparticle. In this study, we found protamine sulfate and heparin increased parental cell labeling^101^, resulting in increased FeEV loading and improved detection when used for MPI. TEM imaging of FeEVs shows that iron is associated with the EV membranes, and proteomics analysis confirms that the FeEVs contain proteins that are considered EV markers. In primary breast tumors *in vivo*, FeEVs from the breast cancer cells were retained for longer and in greater amounts as compared to iron nanoparticles alone, as demonstrated using MPI and confirmed by histological analysis of the tumor. Association with EV membranes also allowed SPIO nanoparticle delivery to the heads of mice when brain metastases were present. MPI could not detect iron in the heads of healthy mice injected with FeEVs, nor in mice with brain metastases injected with SPIO nanoparticles alone. We conclude that association with EV membranes allowed for the cargo, iron nanoparticles, to access metastatic sites across the BBB/BTB. In the future, they could act as a delivery vehicle for therapeutics associated with iron nanoparticles or other therapeutic agents.

## Methods

### Cell Culture⍰

4T1fLuc2 cells (4T1L2; provided by Dr. Bryan Smith, MSU), 4T1BGL cells (provided by Dr. Michael Bachmann, MSU) and 4T1BR5-fLuc/GFP cells (4T1BR5-L2G; provided by Dr. Paula Foster, Western University) were maintained in incubators set at 37°C and 5% CO_2_. Cells were cultured in RPMI+Glutamax with 10% fetal bovine serum (FBS). Cells were counted using the Trypan blue exclusion assay prior to *in vitro* or *in vivo* experiments.

### Iron Labeling of Cells

Three million cells were seeded in a 10 cm^2^ dish for iron labeling. Protamine sulfate (40 µg/ml) or heparin (2 U/ml) and 70 nm dextran-coated Synomag-D (MicroMod, Germany, cat#104-00-701 or cat#126-00-701 (far red fluorescence); 1 mg/ml Fe) were added to 2.5 ml of FBS-free media. Both tubes were well mixed, before the protamine sulfate was added to the heparin and Synomag-D. Five ml of this mixture was added to each plate and 3 to 6-h later 5 ml of complete media was added. The cells were then incubated for 24-h post-addition of iron prior to FeEV isolation (below). The equivalent of 1 dish of FeEVs were used for i.v. biodistribution and primary tumor studies and the equivalent of 2 dishes of FeEVs were used for brain metastasis studies.

### FeEV Isolation Via Differential Centrifugation

Iron-labeled cells were washed 3 times with 10 U/ml heparin and once with PBS, and replaced with media containing 10% EV-depleted FBS. Cells were incubated at 37°C for 24-h to allow for EV production. Conditioned media was centrifuged at 600g for 10 min to remove any cells. The supernatant was then centrifuged at 2,000g for 20 min to remove apoptotic bodies and cell debris. The remaining supernatant containing FeEVs was subsequently centrifuged at 20,000g for 1-h to concentrate FeEVs, which were then washed in PBS and further concentrated via centrifugation at 20,000g for 1-h, before resuspending in PBS.

### EV Isolation via Differential Ultracentrifugation

Non-iron labeled 4T1 cells were seeded at a density of 3×10^6^ cells in a 10 cm dish. Twenty four hours after seeding, the cells were washed twice with PBS to remove traces of media and replaced with media containing 10% EV-depleted FBS. Cells were incubated at 37°C for 24-h to allow for EV production. Conditioned media was centrifuged at 600g for 10 min to remove any cells and the supernatant was centrifuged at 2,000g for 20 min to remove apoptotic bodies and cell debris. Supernatant containing non-labeled EVs was removed and centrifuged at 100,000g for 90 min to concentrate non-labeled EVs. These non-labeled EVs were then washed with PBS and recentrifuged for purity before resuspension in PBS.

### FeEV or SPIO Uptake Into 4T1BR5-L2G Cells in Culture

200,000 4T1BR5-L2G cells were seeded per well, in a 2-well chambered slide (Nunc Lab-Tek II, cat#S6565) and incubated at 37°C and 5% CO_2_ overnight. FeEVs from 3×10^6^ seeded 4T1BR5-L2G cells were collected as above. FeEVs or 20 µg Synomag-D SPIO (far-red fluorescence) were resuspended in PKH26 membrane dye (Sigma-Aldrich, cat#MINI26), as per manufacturers suggestion. FeEVs and SPIO were washed twice with PBS prior to adding to 4T1BR5-L2G cells. Cells and FeEVs or SPIO were incubated at 37°C and 5% CO_2_ for 24-h. Cells were then washed and fixed with 4% paraformaldehyde (PFA). The chamber was removed and cells were coverslipped using Fluoromount-G mounting medium with DAPI (Invitrogen, cat#00-4959-52). Sections were imaged using a Leica DMi8 Thunder microscope equipped with a DFC9000 GTC sCMOS camera and LAS-X software (Leica, Wetzlar, Germany). Large volume computational clearing (LVCC) was performed on the images. Images were prepared using Fiji software^102^.

### Nanoparticle Tracking Analysis

FeEV particle size and concentration were measured using a ZetaView Nanoparticle Tracking Analyzer (Particle Metrix, Germany). An average of 50-150 particles was read per frame as quality control. The analysis parameters used were: Max Area:1000, Min Area: 10, Min Brightness 22, with 11 frames read twice per sample.

### Transmission Electron Microscopy

EVs and FeEVs were visualized via transmission electron microscopy (TEM; JEOL 1400-Flash Transmission Electron Microscope, Japan Electron Optics Laboratory, Japan). Following fixation in 16% PFA, FeEVs were allowed to absorb on 200-mesh, carbon-coated Formvar copper grids for 20 min, before fixation in 2.5% EM-grade glutaraldehyde in 0.1M phosphate buffered saline (PBS) for 15 min at room temperature. The grids were stained with 2% uranyl acetate for contrast and washed with EM-grade PBS and HPLC-grade water before imaging.

### Western Blotting

Cells were lysed in mRIPA lysis buffer (150 mM sodium chloride, 1.0% Triton X-100, 0.25% sodium deoxycholate, 50 mM Tris, pH 7.4) consisting of protease inhibitor (ThermoFisher, A32955) and phosphatase inhibitor (ThermoFisher, A32957). The supernatant was used as cell lysates. Protein concentration of 4T1BR-L2G cell lysate, FeEVs, and EVs isolated using ExoQuick (System Biosciences, CA, USA) was determined using the Pierce BCA Protein Assay kit (ThermoFisher, 23225) using BSA as a standard. The protein quantification curve was completed in replicate 3 times, and the unknowns were all replicated twice.

A total of 15 µg of protein was added per well, mixed with DI H_2_O and RunBlue LDS sample buffer (4X) (Expedeon, NXB31010). The cell lysate mixtures were heated at 70°C for 10 min, while the FeEVs and EVs were not heated to avoid aggregation. The proteins were separated using Mini-PROTEAN TGX Stain-Free Pre-cast gels (BioRad, 4568093) at 100 V for 80-90 min in the BioRad Mini-Protean Tetra system and transferred to a nitrocellulose membrane using the BioRad Trans-Blot Turbo Transfer System, running at 25V for 30 mins. The membrane was then blocked using 5% w/v non-fat dry milk in TBST for one hour at room temperature and then incubated with primary antibody (anti-Alix, 1:5000; Protein Tech, 12422-1-AP or anti-Flotillin-1, 1:5000; Fisher Scientific, BDB610820) at 4°C overnight. The membrane was then washed three times using TBST, and incubated in secondary antibody (Anti-rabbit HRP-linked, 1:2000) at room temperature for one hour. The membrane was washed again three times with TBST, before the Pierce ECL Western Blotting Substrate kit (ThermoFisher, 32209) was added. The proteins and ladder were then imaged using the ChemiDoc MP imaging system (Bio-Rad Laboratories, Inc.) using the autoexposure and chemiluminescence to observe the bands and 635 nm of light with autoexposure for visualization of the ladder.

### LC/MS/MS Analysis

Protein solutions were mixed with 100mM Tris-HCl (pH 8.5) supplemented to 4% (w/v) sodium deoxycholate (SDC) to 270 µl. Samples were reduced and alkylated by adding TCEP and chloroacetamide at 10mM and 40mM, respectively and incubating for 5 min at 45°C with shaking at 2000 rpm in an Eppendorf ThermoMixer C. Trypsin, in 50mM ammonium bicarbonate, was added at a 1:50 ratio (wt/wt) and the mixture was incubated at 37°C overnight with shaking at 1500 rpm in the Thermomixer. Final volume of each digest was ∼300 µl. After digestion, SDC was removed by phase extraction. The samples were acidified to 1% TFA and subjected to C18 solid phase clean up using StageTips1 to remove salts.

An injection of 5 µl (∼600ng) was automatically made using a Thermo (www.thermo.com) EASYnLC 1200 onto a Thermo Acclaim PepMap RSLC 0.1mm x 20mm C18 trapping column and washed for ∼5 min with buffer A. Bound peptides were then eluted over 35 min onto a Thermo Acclaim PepMap RSLC 0.075mm x 500mm resolving column with a gradient of 5%B to 40%B in 24 min, ramping to 90%B at 25 min and held at 90%B for the duration of the run (Buffer A = 99.9% Water/0.1% Formic Acid, Buffer B = 80% Acetonitrile/0.1% Formic Acid/19.9% Water) at a constant flow rate of 300 nl/min. Column temperature was maintained at a constant temperature of 50°C using and integrated column oven (PRSO-V2, Sonation GmbH, Biberach, Germany).

Eluted peptides were sprayed into a ThermoScientific Q-Exactive HF-X mass spectrometer (www.thermo.com) using a FlexSpray spray ion source. Survey scans were taken in the Orbi trap (60000 resolution, determined at m/z 200) and the top 15 ions in each survey scan are then subjected to automatic higher energy collision induced dissociation (HCD) with fragment spectra acquired at 15000 resolution.

Data Analysis was performed as follows. The resulting MS/MS spectra were converted to peak lists using MaxQuant2, v1.6.3.4 (www.maxquant.org), and searched against a protein database containing all mouse sequences available from Uniprot (downloaded from www.uniprot.org, downloaded on 20221114) and appended with common laboratory contaminants using the Andromeda3 search algorithm, a part of the MaxQuant environment. The MaxQuant output was then analyzed using Scaffold, v5.1.2 (www.proteomesoftware.com) to probabilistically validate protein identifications. Assignments validated using the Scaffold 1% FDR confidence filter are considered true.

### Super Resolution Microscopy

Isolated FeEVs from 4T1BR5-L2G cells were analyzed using the ONI EV Profiler kit, a phosphatidylserine-based capture reagent, applied to the EV chip capture surface. The EV sample was then applied and fixed to the surface with ONI EV fixation buffer. Labeled antibodies against tetraspanins (CD81-CF647, CD63-CF568 and CD9-CF488A) were applied to the captured EVs, followed by another fixation step. dSTORM imaging buffer was applied to the samples and imaged on the ONI Nanoimager using dSTORM imaging conditions: 30°C, 52° illumination angle (TIRF), 30 ms exposure per frame. The following lasers were used, sequentially, in a 3000-frame light program: 1000 frames of each 640, 561 and 488 lasers. Analysis was performed using ONI’s cloud-based platform, CODI.

### In Vivo Studies

Six-week old female Balb/C mice were purchased from Charles River Laboratories, and kept in the MSU animal facilities with approval from the MSU Institutional Animal Care and Use Committee.1Mice which did not have tumors received FeEVs (from 4T1BGL; n=2, 1.45×10^10^) or SPIO (Synomag-D far red, n=2, 8.2 μg). Following i.v. administration, the mice were imaged using MPI and CT (described below) at 24-h, 48-h, and 7 d post injection to observe the localization of iron.

Primary tumors were established by injecting 3×10^5^ 4T1L2 cells into the mammary fat pad (MFP) and experiments were initiated 3-weeks after injection. A sample of FeEVs were imaged using MPI prior to injection to determine an estimate of iron present. 4T1L2-derived FeEVs (n=5, 1.7×10^10^) or equal amount SPIO (Synomag-D, n=3, 9.6 μg) were injected into the tumor (intratumoral; i.t.) in 25 μl PBS. Following i.t. administration, mice were imaged with the standard 2D imaging mode in MPI and CT (described below). Following the final imaging time point, mice were sacrificed using 5% carbon dioxide, and underwent post-mortem dissection to remove tumors.

Experimental brain metastasis were initiated in mice via intracardiac (i.c.) injection of 2×10^4^ 4T1BR5-L2G cells, resuspended in 85 µl PBS mixed with 15 µl ultrasound microbubbles (FUJIFILM VisualSonics, WA, USA). Mice were anesthetized (2% isofluorane in oxygen), followed by application of a depilatory to remove fur on their chest. Subcutaneous administration of ketoprofen (5 mg/kg) was used as an analgesic. The mice were placed supine, with their extremities secured, and the left ventricle of the heart was located followed by guidance of the needle and injection of the cell/microbubble mixture using ultrasound (Vevo 2100, Visualsonics). Mice were monitored for brain metastasis (relating to luminescence) using the IVIS Spectrum (PerkinElmer). One hundred microliters of D-Luciferin (PerkinElmer, CT, USA, cat#122799; 30 mg/ml) was injected i.p. 15 min prior to imaging, immediately following i.c. cell injection and every 48-h until FeEV administration. A sample of FeEVs were imaged using MPI prior to injection to determine an estimate of iron present. 4T1BR5-L2G-derived FeEVs (6.97×10^10^) or SPIO (30 or 36 µg) were injected i.c. 7-8 days following brain metastasis establishment. Mice were given analgesic (ketoprofen, 5 mg/kg) subcutaneous prior to beginning the procedure. Ultrasound was used (as above) to administer FeEVs or SPIOs. Following i.c. injection, mice were imaged using MPI and CT (described below).

### In Vivo Imaging

Imaging was performed at the time-points described, using the following parameters. MPI was performed using the standard 3D and 2D imaging modes (FOV 12×6×6 cm, 5.7 T/m gradient, 1 (2D) or 21 (3D) projections, and 1 average). Mice were then transferred to a Quantum GX microCT scanner (PerkinElmer). Whole-body CT images were acquired using 3×8 s scans with the following parameters: 90 kV voltage, 88 μA amperage, 72 mm acquisition FOV, and 60 mm reconstruction FOV, resulting in 240 μm voxels Standards of known iron amount were placed in the MPI bed to aid in co-registration of μCT and MPI scans.

### Image Analysis

MPI data sets were visualized and analyzed using Horos imaging software (Horos is a free and open source code software program that is distributed free of charge under the LGPL license at Horosproject.org and sponsored by Nimble Co LLC d/b/a Purview in Annapolis, MD, USA).

MPI quantification was performed on 2D images. Signal threshold was chosen by selecting an area of background from the image which did not have any iron present (primary tumor: adjacent gut signal and brain: signal outside of the head). 3x sdev of the background was set as a lower threshold to capture signal above this value using thresholding in Horos. For primary tumor quantification, the mean signal of the gut (minus the noise from a blank image) was subtracted from the mean signal from the tumor to account for any signal which was added to the tumor. In instances where the signal was low (*i.e*., SPIO injection, 72-h), if the thresholding spread to include other regions (*i.e*., the iron fiducials), it was removed manually. Iron signal in FeEV pellets, the primary tumor, or the brain of mice after injection of FeEV, EV or SPIO were determined as above, with total MPI signal calculated by mean signal x area. Mass of iron in the FeEV samples and measurements from NTA (above) were used to determine the amount of iron per FeEV.

Iron amount was determined using calibration lines. Different amounts of iron were imaged with MPI using imaging sequences to establish a reference curve for each scan type utilized: standard and high sensitivity 2D or standard 3D. A simple linear regression was performed to find the slope of the data (m) using best-fit values (y=known iron content and x= MPI signal) with the x,y intercept set to 0. The equation y=mx allowed for quantification of the iron content in FeEV pellets and *in vivo* by substituting the total MPI signal (x) from the ROI into the equation to solve for iron (y).

Iron concentration values displayed on scale bars were determined by plotting mean signal (x) and iron/mm^2^ or iron/mm^3^ (y, based on 2D or 3D data sets) from the iron amounts used for the calibration lines. Concentration of iron was solved for by inputting mean signal (x) and solving for y.

### Histology

Brains and isolated primary tumors were fixed overnight in 4% paraformaldehyde followed by cryopreservation through serial submersion in 10%, 20%, and 30% sucrose for 24-h each. Samples were then placed in a bed of OCT, before being flash frozen in a mixture of dry ice and ethanol. The frozen samples were stored at -20°C in preparation for tissue sectioning using a cryostat (Leica CM3050 S, 10 µm thickness).

For primary tumors, tissue sections with far-red expression from the iron nanoparticles, as determined by screening, were washed with PBS for 5 min, before being added to 0.3% triton X-100 in PBS and incubated for 45 min. Slides were then incubated in blocking buffer (5% goat serum and 0.3% triton X-100 in PBS) for 60 min. Anti-CD47-PE (3 µg/ml; Biolegend, cat#127507) and F4/80 Monoclonal Antibody (1:200; ThermoFisher, cat#14-480185) were then added to the slides overnight at 4°C followed by washing with PBS. Sections were then incubated with a secondary Goat anti-Rat IgG antibody, AF647 (1:500, ThermoFisher, cat#A-21247) for 2 hours at room temperature followed by washing with PBS. Sections were imaged using a Leica DMi8 Thunder microscope equipped with a DFC9000 GTC sCMOS camera and LAS-X software (Leica, Wetzlar, Germany). Large volume computational clearing (LVCC) was performed on the images. Images were prepared using Fiji software^102^.

Brain sections were washed with PBS for 5 min followed by Perls’ Prussian Blue (PPB) staining to visualize iron. DAPI mounting media (Fluoromount-G, Invitrogen) was used to visualize nuclei. Sections were imaged using a Nikon Eclipse Ci microscope equipped with a Nikon DS-Fi3 high-definition camera (Nikon Instruments Inc. Tokyo, Japan) for color and brightfield acquisition, CoolSNAP DYNO (Photometrics, AZ, USA) for fluorescent imaging and NIS elements BR 5.21.02 software (Nikon). Images were prepared using Fiji software^102^. PPB staining (blue) was pseudocolored magenta for overlay with DAPI.

### Statistical Analysis

Statistical analyses were performed using Prism software (10.1.1, GraphPad Inc., CA, USA). A two-way repeated measures ANOVA with uncorrected Fisher’s LSD was used to compare differences in EV associated proteins between FeEVs and EVs derived from 4T1L2 or 4T1BR5-L2G cells. A two-way repeated measures ANOVA with uncorrected Fisher’s LSD was used to compare differences in iron quantification in primary tumors between those with that received FeEVs or SPIO, and between 0, 24, 48 and 72-h. Data are expressed as mean +/-standard deviation; *p*<0.05 was considered a significant finding.

## Supporting Information

Representative FeEV characteristics and iron content for FeEVs injected *in vivo*, MPI of Synomag-D SPIO and FeEV pellet, western blot analysis of FeEV, EV and cell lysate for 4T1BR5-L2G, super resolution microscopy of FeEVs, *in vivo* biodistribution of SPIO or FeEVs into healthy mice.

## Acknowledgements

We would like to thank Drs. Michael Bachmann, Bryan Smith and Paula Foster for providing us with the cells used. We acknowledge the MSU Mass Spectrometry and Metabolomics Core for performing the experiments and providing the protocol that was used for the analysis. We thank the MSU IQ Advanced Molecular Imaging Facility for their help and guidance during animal imaging. We acknowledge the MSU Center for Advanced Microscopy for the use of their protocols, equipment, and facilities. We thank Dr. Kanada for sharing his expertise with EVs. We thank the applications team at ONI for their support with the super resolution microscopy. This study was funded by METAvivor (AVM) and the James and Kathleen Cornelius Endowment (CHC). TOC graphic was made with BioRender.

## Notes

### Competing Interest Statement

The authors have declared no competing interest.

### Summary of Updates

Added figure 2 - in vitro uptake Added supplementary figure 1 - MPI of SPIO and FeEV pellet. Figure 3 - included 0, 24 and 72h time points of iron in tumors

## References

1. Ferlay, J. et al. Global Cancer Observatory: Cancer Today. (2024).

2. Sun, H., Xu, J., Dai, S., Ma, Y. & Sun, T. Breast cancer brain metastasis: Current evidence and future directions. Cancer Med 12, 1007–1024 (2023).

3. Farahani, M. K., Gharibshahian, M., Rezvani, A. & Vaez, A. Breast cancer brain metastasis: from etiology to state-of-the-art modeling. J Biol Eng 17, 41 (2023).

4. Dubey, A., Agrawal, S., Agrawal, V., Dubey, T. & Jaiswal, A. Breast Cancer and the Brain: A Comprehensive Review of Neurological Complications. Cureus 15, e48941 (2023).

5. Bansal, R., Van Swearingen, A. E. D. & Anders, C. K. Triple Negative Breast Cancer and Brain Metastases. Clin Breast Cancer 23, 825–831 (2023).

6. Ivanova, M. et al. Breast Cancer with Brain Metastasis: Molecular Insights and Clinical Management. Genes (Basel) 14, 1160 (2023).

7. Rader, R. K., Anders, C. K., Lin, N. U. & Sammons, S. L. Available Systemic Treatments and Emerging Therapies for Breast Cancer Brain Metastases. Curr Treat Options Oncol 24, 611–627 (2023).

8. Nieder, C., Andratschke, N. H. & Grosu, A. L. Brain Metastases: Is There Still a Role for Whole-Brain Radiation Therapy? Semin Radiat Oncol 33, 129–138 (2023).

9. Zhao, H., Wang, L., Ji, X., Zhang, L. & Li, C. Biology of breast cancer brain metastases and novel therapies targeting the blood brain barrier: an updated review. Medical Oncology 40, 181 (2023).

10. Steeg, P. S. The blood–tumour barrier in cancer biology and therapy. Nat Rev Clin Oncol 18, 696–714 (2021).

11. Steeg, P. S. The blood–tumour barrier in cancer biology and therapy. Nat Rev Clin Oncol 18, 696–714 (2021).

12. Adkins, C. E. et al. Characterization of passive permeability at the blood–tumor barrier in five preclinical models of brain metastases of breast cancer. Clin Exp Metastasis 33, 373–383 (2016).

13. Lockman, P. R. et al. Heterogeneous blood–tumor barrier permeability determines drug efficacy in experimental brain metastases of breast cancer. Clinical cancer research 16, 5664–5678 (2010).

14. Sakamoto, Y., Ochiya, T. & Yoshioka, Y. Extracellular vesicles in the breast cancer brain metastasis: physiological functions and clinical applications. Front Hum Neurosci 17, 1278501 (2023).

15. Luo, T., Kang, Y., Liu, Y., Li, J. & Li, J. Small extracellular vesicles in breast cancer brain metastasis and the prospect of clinical application. Front Bioeng Biotechnol 11, 1162089 (2023).

16. Welsh, J. A. et al. Minimal information for studies of extracellular vesicles (MISEV2023): From basic to advanced approaches. J Extracell Vesicles 13, e12404 (2024).

17. Chen, C. C. et al. Elucidation of exosome migration across the blood–brain barrier model in vitro. Cell Mol Bioeng 9, 509–529 (2016).

18. Morad, G. et al. Tumor-derived extracellular vesicles breach the intact blood–brain barrier via transcytosis. ACS Nano 13, 13853–13865 (2019).

19. Zhuang, X. et al. Treatment of brain inflammatory diseases by delivering exosome encapsulated anti-inflammatory drugs from the nasal region to the brain. Molecular Therapy 19, 1769–1779 (2011).

20. Alvarez-Erviti, L. et al. Delivery of siRNA to the mouse brain by systemic injection of targeted exosomes. Nat Biotechnol 29, 341–345 (2011).

21. Betzer, O. et al. In Vivo Neuroimaging of Exosomes Using Gold Nanoparticles. ACS Nano 11, 10883-10893–10893 (2017).

22. Zhuang, X. et al. Treatment of brain inflammatory diseases by delivering exosome encapsulated anti-inflammatory drugs from the nasal region to the brain. Molecular Therapy 19, 1769–1779 (2011).

23. Tian, Y. et al. A doxorubicin delivery platform using engineered natural membrane vesicle exosomes for targeted tumor therapy. Biomaterials 35, 2383–2390 (2014).

24. Srivastava, A. et al. Nanosomes carrying doxorubicin exhibit potent anticancer activity against human lung cancer cells. Sci Rep 6, 38541 (2016).

25. Kim, M. S. et al. Development of exosome-encapsulated paclitaxel to overcome MDR in cancer cells. Nanomedicine 12, 655–664 (2016).

26. Han, Z. et al. Highly efficient magnetic labelling allows MRI tracking of the homing of stem cell-derived extracellular vesicles following systemic delivery. J Extracell Vesicles 10, e12054 (2021).

27. Jia, G. et al. NRP-1 targeted and cargo-loaded exosomes facilitate simultaneous imaging and therapy of glioma in vitro and in vivo. Biomaterials 178, 302–316 (2018).

28. Dabrowska, S. et al. Imaging of extracellular vesicles derived from human bone marrow mesenchymal stem cells using fluorescent and magnetic labels. Int J Nanomedicine 1653–1664 (2018).

29. Han, Z. et al. Highly efficient magnetic labelling allows MRI tracking of the homing of stem cell-derived extracellular vesicles following systemic delivery. J Extracell Vesicles 10, e12054 (2021).

30. Kim, H. Y. et al. Mesenchymal stem cell-derived magnetic extracellular nanovesicles for targeting and treatment of ischemic stroke. Biomaterials 243, 119942 (2020).

31. Lara, P. et al. Gold nanoparticle based double-labeling of melanoma extracellular vesicles to determine the specificity of uptake by cells and preferential accumulation in small metastatic lung tumors. J Nanobiotechnology 18, 1–17 (2020).

32. Hu, L., Wickline, S. A. & Hood, J. L. Magnetic resonance imaging of melanoma exosomes in lymph nodes. Magn Reson Med 74, 266–271 (2015).

33. Zhang, H. et al. Iron Oxide Nanoparticles Engineered Macrophage-Derived Exosomes for Targeted Pathological Angiogenesis Therapy. ACS Nano (2024) doi:10.1021/acsnano.4c00699.

34. Jung, K. O., Jo, H., Yu, J. H., Gambhir, S. S. & Pratx, G. Development and MPI tracking of novel hypoxia-targeted theranostic exosomes. Biomaterials 177, 139–148 (2018).

35. Nguyen, V. Du, Kim, H. Y., Choi, Y. H., Park, J.-O. & Choi, E. Tumor-derived extracellular vesicles for the active targeting and effective treatment of colorectal tumors in vivo. Drug Deliv 29, 2621–2631 (2022).

36. Qiao, L. et al. Tumor cell-derived exosomes home to their cells of origin and can be used as Trojan horses to deliver cancer drugs. Theranostics 10, 3474 (2020).

37. Lin, S.-W., Tsai, J.-C. & Shyong, Y.-J. Drug delivery of extracellular vesicles: Preparation, delivery strategies and applications. Int J Pharm 642, 123185 (2023).

38. Hill, M. L. et al. Exosome-Coated Prussian Blue Nanoparticles for Specific Targeting and Treatment of Glioblastoma. ACS Appl Mater Interfaces (2024) doi:10.1021/acsami.4c02364.

39. Yu, E. Y. et al. Magnetic Particle Imaging: A Novel in Vivo Imaging Platform for Cancer Detection. Nano Lett 17, 1648–1654 (2017).

40. Panagiotopoulos, N. et al. Magnetic particle imaging: current developments and future directions. Int J Nanomedicine 10, 3097–3114 (2015).

41. Bulte, J. W. M. Superparamagnetic iron oxides as MPI tracers: A primer and review of early applications. Adv Drug Deliv Rev 138, 293–301 (2019).

42. Williams, R. J. et al. Dual magnetic particle imaging and akaluc bioluminescence imaging for tracking cancer cell metastasis. Tomography 9, 178–194 (2023).

43. Huang, X. et al. Deep penetrating and sensitive targeted magnetic particle imaging and photothermal therapy of early-stage glioblastoma based on a biomimetic nanoplatform. Advanced Science 10, 2300854 (2023).

44. Makela, A. V et al. Magnetic Particle Imaging of Macrophages Associated with Cancer: Filling the Voids Left by Iron-Based Magnetic Resonance Imaging. Mol Imaging Biol 22, 958–968 (2020).

45. Makela, A. V et al. Tracking the fates of iron-labeled tumor cells in vivo using Magnetic Particle Imaging. bioRxiv 2021.10.06.463387 (2021) doi:10.1101/2021.10.06.463387.

46. Szwargulski, P. et al. Monitoring Intracranial Cerebral Hemorrhage Using Multicontrast Real-Time Magnetic Particle Imaging. ACS Nano 14, 13913–13923 (2020).

47. Ludewig, P. et al. Magnetic Particle Imaging for Real-Time Perfusion Imaging in Acute Stroke. ACS Nano 11, 10480–10488 (2017).

48. Graeser, M. et al. Design of a head coil for high resolution mouse brain perfusion imaging using magnetic particle imaging. Phys Med Biol 65, 235007 (2020).

49. Cooley, C. Z., Mandeville, J. B., Mason, E. E., Mandeville, E. T. & Wald, L. L. Rodent Cerebral Blood Volume (CBV) changes during hypercapnia observed using Magnetic Particle Imaging (MPI) detection. Neuroimage 178, 713–720 (2018).

50. Yu, E. Y. et al. Magnetic Particle Imaging for Highly Sensitive, Quantitative, and Safe in Vivo Gut Bleed Detection in a Murine Model. ACS Nano 11, 12067–12076 (2017).

51. Barry Fung, K. L. et al. Rapid in situ labelling and tracking of neutrophils and macrophages to inflammation using antibody-functionalized MPI tracers. Int J Magn Part Imaging 8, (2022).

52. Chandrasekharan, P. et al. Non-radioactive and sensitive tracking of neutrophils towards inflammation using antibody functionalized magnetic particle imaging tracers. Nanotheranostics 5, 240 (2021).

53. Zhu, X., Li, J., Peng, P., Hosseini-nassab, N. & Smith, B. R. Quantitative drug release monitoring in tumors of living subjects by magnetic particle imaging nanocomposite Quantitative drug release monitoring in tumors of living subjects by magnetic particle imaging nanocomposite. Nano Lett (2019) doi:10.1021/acs.nanolett.9b01202.

54. Kuo, R., Chandrasekharan, P., Fung, K. L. B. & Conolly, S. In vivo therapeutic cell tracking using magnetic particle imaging. Int J Magn Part Imaging 8, (2022).

55. Sehl, O. C., Makela, A. V, Hamilton, A. M. & Foster, P. J. Trimodal Cell Tracking In Vivo: Combining Iron- and Fluorine-Based Magnetic Resonance Imaging with Magnetic Particle Imaging to Monitor the Delivery of Mesenchymal Stem Cells and the Ensuing Inflammation. Tomography 5, 367–376 (2019).

56. Wang, P. et al. Magnetic particle imaging of islet transplantation in the liver and under the kidney capsule in mouse models. Quant Imaging Med Surg 8, 114 (2018).

57. Bulte, J. W. M. et al. Quantitative ‘Hot Spot’ Imaging of Transplanted Stem Cells using Superparamagnetic Tracers and Magnetic Particle Imaging. Tomography 1, 91–97 (2015).

58. Zheng, B. et al. Magnetic particle imaging tracks the long-term fate of in vivo neural cell implants with high image contrast. Sci Rep 5, 1–9 (2015).

59. Zheng, B. et al. Quantitative magnetic particle imaging monitors the transplantation, biodistribution, and clearance of stem cells in vivo. Theranostics 6, 291–301 (2016).

60. Song, G. et al. Janus Iron Oxides @ Semiconducting Polymer Nanoparticle Tracer for Cell Tracking by Magnetic Particle Imaging. Nano Lett 18, 182–189 (2018).

61. Avugadda, S. K. et al. Uncovering the magnetic particle imaging and magnetic resonance imaging features of iron oxide nanocube clusters. Nanomaterials 11, 1–11 (2021).

62. Gloag, L. et al. Zero valent iron core–iron oxide shell nanoparticles as small magnetic particle imaging tracers. Chemical communications 56, 3504–3507 (2020).

63. Fung, K. L. B. et al. First superferromagnetic remanence characterization and scan optimization for super-resolution Magnetic Particle Imaging. Nano Lett 23, 1717–1725 (2023).

64. Wang, Q. et al. Artificially Engineered Cubic Iron Oxide Nanoparticle as a High-Performance Magnetic Particle Imaging Tracer for Stem Cell Tracking. ACS Nano 14, 2053–2062 (2020).

65. Tay, Z. W. et al. Superferromagnetic nanoparticles enable order-of-magnitude resolution & sensitivity gain in magnetic particle imaging. Small Methods 5, 2100796 (2021).

66. Lu, K., Goodwill, P., Zheng, B. & Conolly, S. Multi-channel acquisition for isotropic resolution in magnetic particle imaging. IEEE Trans Med Imaging 37, 1989–1998 (2017).

67. Pagan, J., McDonough, C., Vo, T. & Tonyushkin, A. Single-sided magnetic particle imaging device with field-free-line geometry for in vivo imaging applications. IEEE Trans Magn 57, 1–5 (2020).

68. Hayat, H. et al. Artificial intelligence analysis of magnetic particle imaging for islet transplantation in a mouse model. Mol Imaging Biol 23, 18–29 (2021).

69. Sehl, O. C. et al. MPI region of interest (ROI) analysis and quantification of iron in different volumes. 5–7 (2022).

70. Liu, H. et al. Cancer stem cells from human breast tumors are involved in spontaneous metastases in orthotopic mouse models. Proceedings of the National Academy of Sciences 107, 18115–18120 (2010).

71. Welsh, J. A. et al. Minimal information for studies of extracellular vesicles (MISEV2023): From basic to advanced approaches. J Extracell Vesicles 13, e12404 (2024).

72. May, R. C. & Machesky, L. M. Phagocytosis and the actin cytoskeleton. J Cell Sci 114, 1061–1077 (2001).

73. Garin, J. et al. The Phagosome Proteome: Insight into Phagosome Functions. The Journal of Cell Biology vol. 152 http://www.jcb.org/cgi/content/full/152/1/165 (2001).

74. Yanatori, I., Richardson, D. R., Dhekne, H. S., Toyokuni, S. & Kishi, F. CD63 is regulated by iron via the IRE-IRP system and is important for ferritin secretion by extracellular vesicles. Blood 138, 1490–1503 (2021).

75. Andreu, Z. & Yáñez-Mó, M. Tetraspanins in extracellular vesicle formation and function. Front Immunol 5, 109543 (2014).

76. Jankovičová, J., Sečová, P., Michalková, K. & Antalíková, J. Tetraspanins, more than markers of extracellular vesicles in reproduction. Int J Mol Sci 21, 7568 (2020).

77. Tognoli, M. L. et al. Lack of involvement of CD63 and CD9 tetraspanins in the extracellular vesicle content delivery process. Commun Biol 6, 532 (2023).

78. Glebov, O. O., Bright, N. A. & Nichols, B. J. Flotillin-1 defines a clathrin-independent endocytic pathway in mammalian cells. Nat Cell Biol 8, 46–54 (2006).

79. Fanaei, M., Monk, P. N. & Partridge, L. J. The role of tetraspanins in fusion. Biochem Soc Trans 39, 524–528 (2011).

80. Liu, Y.-J. & Wang, C. A review of the regulatory mechanisms of extracellular vesicles-mediated intercellular communication. Cell Communication and Signaling 21, 77 (2023).

81. Rana, S., Yue, S., Stadel, D. & Zöller, M. Toward tailored exosomes: The exosomal tetraspanin web contributes to target cell selection. Int J Biochem Cell Biol 44, 1574–1584 (2012).

82. Xie, F. et al. Breast cancer cell-derived extracellular vesicles promote CD8+1T cell exhaustion via TGF-β type II receptor signaling. Nat Commun 13, 4461 (2022).

83. Gerwing, M. et al. Tracking of Tumor Cell–Derived Extracellular Vesicles In Vivo Reveals a Specific Distribution Pattern with Consecutive Biological Effects on Target Sites of Metastasis. Mol Imaging Biol 22, 1501–1510 (2020).

84. Frolova, L. & Li, I. T. S. Targeting capabilities of native and bioengineered extracellular vesicles for drug delivery. Bioengineering 9, 496 (2022).

85. Lázaro-Ibáñez, E. et al. Metastatic state of parent cells influences the uptake and functionality of prostate cancer cell-derived extracellular vesicles. J Extracell Vesicles 6, 1354645 (2017).

86. Park, J., Lee, H., Youn, Y. S., Oh, K. T. & Lee, E. S. Tumor-homing pH-sensitive extracellular vesicles for targeting heterogeneous tumors. Pharmaceutics 12, 372 (2020).

87. Liu, Y. et al. Focused ultrasound-augmented targeting delivery of nanosonosensitizers from homogenous exosomes for enhanced sonodynamic cancer therapy. Theranostics 9, 5261 (2019).

88. Makela, A. V et al. Magnetic Particle Imaging of Macrophages Associated with Cancer: Filling the Voids Left by Iron-Based Magnetic Resonance Imaging. Mol Imaging Biol 22, 958–968 (2020).

89. Vermaelen, K. & Pauwels, R. Accurate and simple discrimination of mouse pulmonary dendritic cell and macrophage populations by flow cytometry: methodology and new insights. Cytometry A 61, 170–177 (2004).

90. Makela, A. V, Gaudet, J. M. & Foster, P. J. Quantifying tumor associated macrophages in breast cancer: a comparison of iron and fluorine-based MRI cell tracking. Sci Rep 7, (2017).

91. Makela, A. V & Foster, P. J. Imaging macrophage distribution and density in mammary tumors and lung metastases using fluorine-19 MRI cell tracking. Magn Reson Med 80, 1138–1147 (2018).

92. Daldrup-Link, H. E. et al. MRI of tumor-associated macrophages with clinically applicable iron oxide nanoparticles. Clin Cancer Res 17, 5695–5704 (2011).

93. Kodali, M. et al. A single intranasal dose of human mesenchymal stem cell-derived extracellular vesicles after traumatic brain injury eases neurogenesis decline, synapse loss, and BDNF-ERK-CREB signaling. Front Mol Neurosci 16, 1185883 (2023).

94. Zhou, X. et al. Intranasal delivery of BDNF-loaded small extracellular vesicles for cerebral ischemia therapy. Journal of Controlled Release 357, 1–19 (2023).

95. Li, J. et al. The landscape of extracellular vesicles combined with intranasal delivery towards brain diseases. Nano Today 55, 102169 (2024).

96. Zhuo, Z. et al. Targeted extracellular vesicle delivery systems employing superparamagnetic iron oxide nanoparticles. Acta Biomater 134, 13–31 (2021).

97. Lee, J.-R. et al. Nanovesicles derived from iron oxide nanoparticles–incorporated mesenchymal stem cells for cardiac repair. Sci Adv 6, eaaz0952 (2024).

98. Abdelsalam, M., Ahmed, M., Osaid, Z., Hamoudi, R. & Harati, R. Insights into Exosome Transport through the Blood–Brain Barrier and the Potential Therapeutical Applications in Brain Diseases. Pharmaceuticals 16, 571 (2023).

99. Salimi, L. et al. Physiological and pathological consequences of exosomes at the blood–brain-barrier interface. Cell Communication and Signaling 21, 1–22 (2023).

100. Fitzgerald, D. P. et al. Reactive glia are recruited by highly proliferative brain metastases of breast cancer and promote tumor cell colonization. Clin Exp Metastasis 25, 799–810 (2008).

101. Arbab, A. S. et al. Efficient magnetic cell labeling with protamine sulfate complexed to ferumoxides for cellular MRI. Blood 104, 1217–1224 (2004).

102. Schindelin, J. et al. Fiji: an open-source platform for biological-image analysis. Nat Methods 9, 676–682 (2012).

