## Supplementary material for "Magnetic particle imaging reveals that iron-labeled extracellular vesicles accumulate in brains of mice with metastases": Table S1

| | Average peak size | Number of FeEVs administered | Total iron administered (ug) | Iron (ug)/EV | Seeded cell density |
|---|---|---|---|---|---|
| Healthy + 4T1BGL FeEVs | 93.1 nm (n=3) | $1.45x10^{10}$ | 6.13 ug | $4.24x10^{-10}$ | $3x10^{6}$ |
| Primary tumors + 4T1L2 FeEVs | 95.6 nm (n=3) | $1.70x10^{10}$ | 9.63 ug | $5.66x10^{-10}$ | $3x10^{6}$ |
| Brain metastasis + 4T1BR5-L2G FeEVs | 101.1 nm (n=3) | $6.97x10^{10}$ | 35.63 ug | $5.11x10^{-10}$ | $6x10^{6}$ |

**Table S1**. Representative FeEV characteristics and iron content for FeEVs injected into non-tumor bearing (healthy) mice, mice with primary tumors and mice with brain metastasis.
