## Supplementary material for "Magnetic particle imaging reveals that iron-labeled extracellular vesicles accumulate in brains of mice with metastases": Figure S1

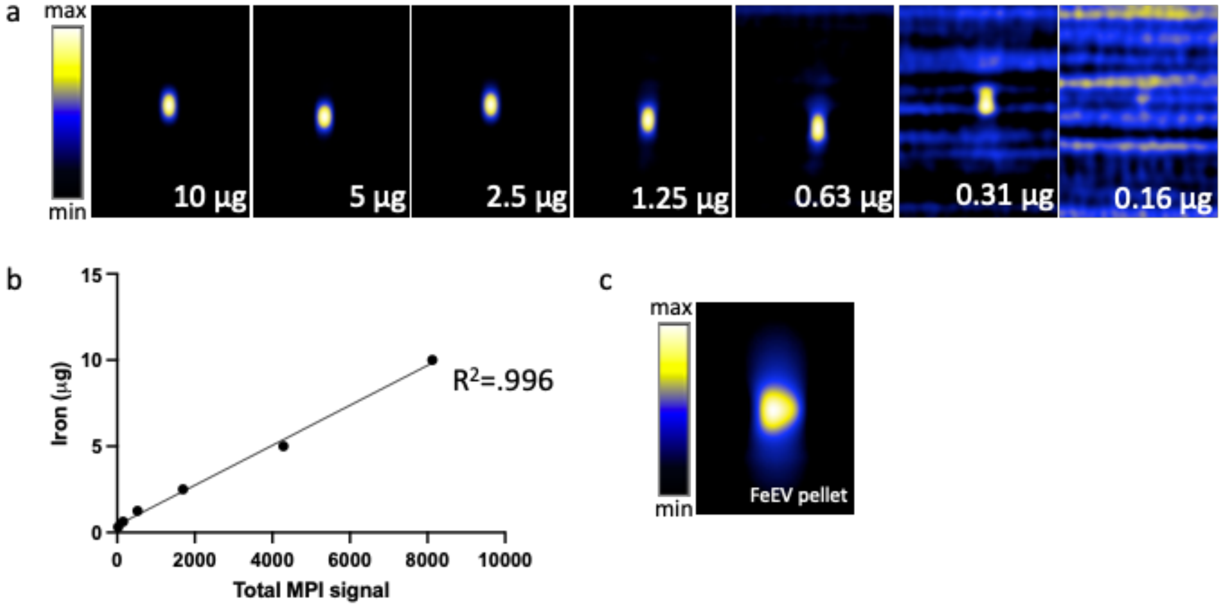


**Figure S1**. Magnetic Particle Imaging of Synomag-D SPIO and FeEV pellet. Different amounts of Synomag-D (0.16 μg – 10 μg) in 1 μl volumes imaged by MPI (**a**). There is a linear relationship between known iron amount (μg, y) and total MPI signal (x) (**b**). Representative FeEV pellet imaged by MPI prior to *in vivo* administration (**c**).
