## Supplementary material for "Magnetic particle imaging reveals that iron-labeled extracellular vesicles accumulate in brains of mice with metastases": Figure S2

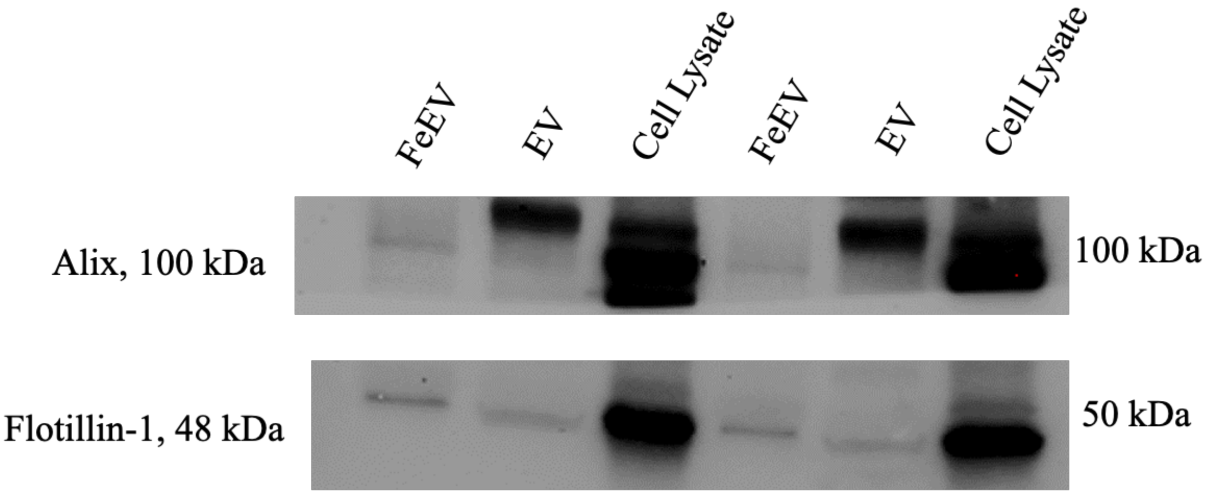


**Figure S2**. Western blot analysis of 4T1BR5-L2G-derived iron-labeled EVs (FeEVs), EVs and cell lysate. FeEVs, EVs and cell lysate from 4T1BR5-L2G cells contain Alix (upper row) and Flotillin-1 (lower row). Repeated data is a technical replicate.
