## Supplementary material for "Magnetic particle imaging reveals that iron-labeled extracellular vesicles accumulate in brains of mice with metastases": Figure S3

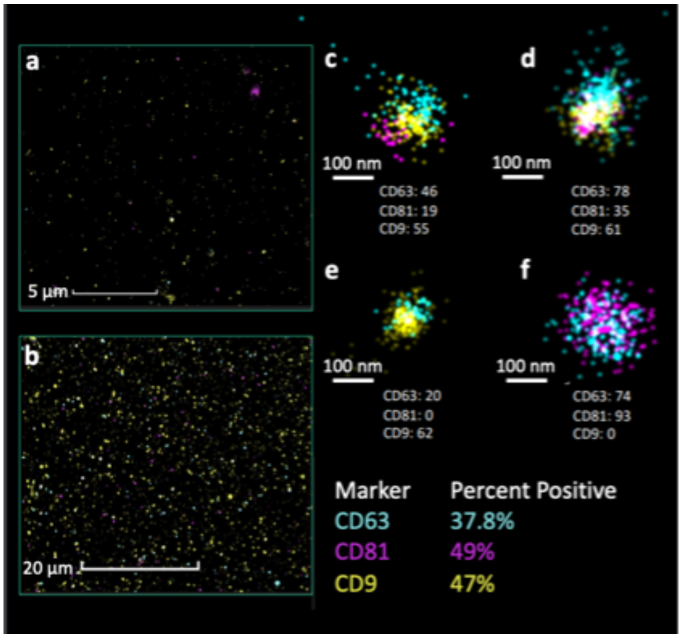


**Figure S3**. Super resolution microscopy of iron-labeled extracellular vesicles (FeEVs) derived from 4T1BR5-L2G cells. The sample was surface stained with anti-CD63 (CF568, cyan), anti-CD81 (CF647, magenta) and anti-CD9 (CF488A, yellow) antibodies. dSTORM imaging of sample at different magnifications (**a,b**). Four different FeEVs are shown (**c-f**), with different compositions of markers.
