## Supplementary material for "Magnetic particle imaging reveals that iron-labeled extracellular vesicles accumulate in brains of mice with metastases": Figure S4

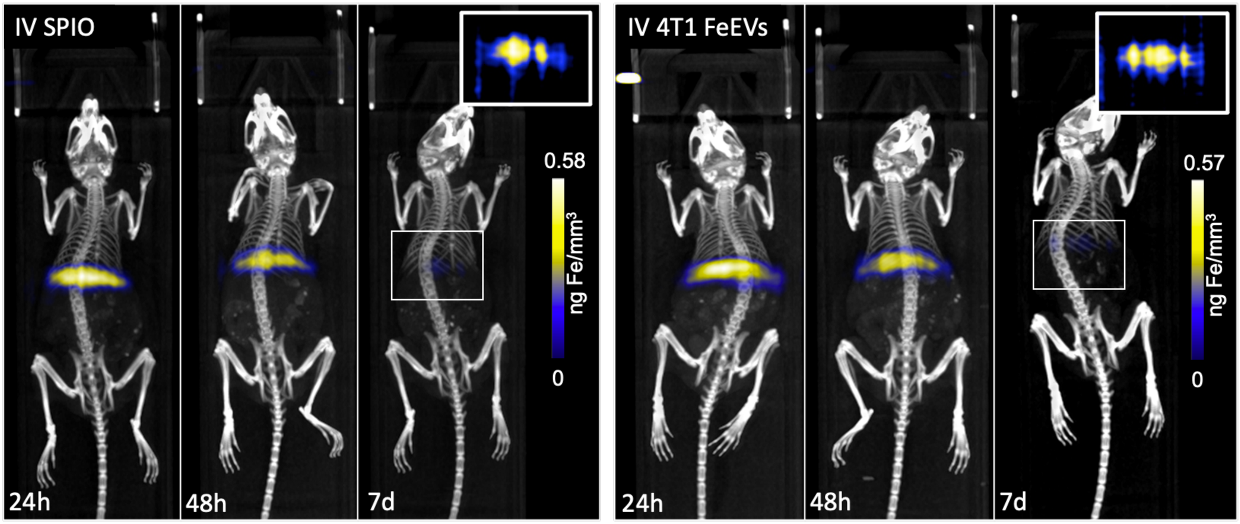


**Figure S4**. *In vivo* biodistribution of SPIO or 4T1 iron-labeled EVs (FeEVs) in non-tumor bearing mice. SPIO (left) or 4T1-derived FeEVs (right) were administered intravenous into non-tumor bearing mice. Magnetic particle imaging (MPI) and CT were performed at 24-hours (h), 48-h and 7-days (d); images are overlays of the MPI and CT images with intensity scale consistent longitudinally. Inset, top right is full dynamic range of signal from the liver at 7-d post injection.
